# Simulations suggest a constrictive force is required for Gram-negative bacterial cell division

**DOI:** 10.1101/406389

**Authors:** Lam T. Nguyen, Catherine M. Oikonomou, H. Jane Ding, Mohammed Kaplan, Qing Yao, Yi-Wei Chang, Morgan Beeby, Grant J. Jensen

## Abstract

To divide, Gram-negative bacterial cells must remodel their peptidoglycan cell wall to a smaller and smaller radius at the division site, but how this process occurs remains debated. While the tubulin homolog FtsZ is thought to generate a constrictive force, it has also been proposed that cell wall remodeling alone is sufficient to drive membrane constriction, possibly via a make-before-break mechanism in which new hoops of cell wall are made inside the existing hoops (make) before bonds in the existing wall are cleaved (break). Previously, we constructed software, REMODELER 1, to simulate cell wall remodeling in rod-shaped bacteria during growth. Here, we used this software as the basis for an expanded simulation system, REMODELER 2, which we used to explore different mechanistic models of cell wall division. We found that simply organizing the cell wall synthesis complexes at the midcell was not sufficient to cause wall invagination, even with the implementation of a make-before-break mechanism. Applying a constrictive force at the midcell could drive division if the force was sufficiently large to initially constrict the midcell into a compressed state before new hoops of relaxed cell wall were incorporated between existing hoops. Adding a make-before-break mechanism could drive division with a smaller constrictive force sufficient to bring the midcell peptidoglycan into a relaxed, but not necessarily compressed, state.

## Introduction

Bacterial cells are protected from turgor pressure by a peptidoglycan (PG) cell wall that is composed of long glycan strands crosslinked by short peptides (Vollmer et al., 2008). This relatively rigid sacculus allows cells to adopt specialized shapes, such as the rod shape of many Gram-negative bacteria. In order for the cell to change size or shape during growth and division, the pressurized sacculus must be carefully remodeled. This is accomplished by a set of cell wall remodeling enzymes including transglycosylases, transpeptidases, and endopeptidases. Experimental insights into the exact molecular mechanisms of these remodeling enzymes and how their functions are coordinated remain limited. Previously, we gained insight into these questions by building simulation software, REMODELER 1, to study cell wall synthesis during cell elongation (Nguyen et al., 2015). In this software, a cylindrical cell wall is coarse-grained as chains of tetrasaccharide beads running circumferentially around the cylinder and connected by peptide crosslinks. The functions of transglycosylases, transpeptidases, and endopeptidases are explicitly modeled as beads. Using this software, we found that in order to maintain the integrity and rod shape of the cell, these remodeling enzymes have to coordinate with one another locally in synthetic complexes, but that no long-range coordination of the independent complexes is required. We also found that these complexes must contain a lytic transglycosylase to remove long, uncrosslinked glycan tails to clear the path for enzyme movement (Nguyen et al., 2015). (Such an enzyme was independently identified experimentally (Cho et al., 2014).)

During cell elongation, the diameter of a rod-shaped cell is conserved. In contrast, during division, the diameter of the cell wall at the division site must become smaller and smaller. How the cell overcomes turgor pressure to remodel its cell wall to a smaller diameter remains unclear (Osawa and Erickson, 2018). It is unlikely to be due to a fundamentally different mode of synthesis, since (a) partially overlapping and homologous sets of enzymes mediate remodeling in cell growth and division (Egan and Vollmer, 2012); (b) these PG synthesis enzymes were shown to move around the cell’s circumference during both elongation (Domínguez-Escobar et al., 2011; Garner et al., 2011) and division (Bisson-Filho et al., 2017; Yang et al., 2017); and (c) in purified sacculi, glycan strands exhibit similar circumferential orientation throughout the length of the cell (Gan et al., 2008; Turner et al., 2018).

The protein FtsZ, a tubulin homolog found in nearly all bacteria and many archaea, forms filaments at the midcell during cell division (Bi and Lutkenhaus, 1991; Li et al., 2007; Szwedziak et al., 2014; Yao et al., 2017). It has been proposed that these filaments exert a constrictive force on the membrane and serve as a scaffold for the cell wall synthesis machinery (Erickson et al., 2010). Based on cryo-electron microscopy images of dividing cells, it has been proposed that GTP-hydrolyzing FtsZ filaments can generate a constrictive force either by switching conformation from straight to curved (Li et al., 2007) or by overlapping to form a closed ring which then tightens to constrict the membrane (Szwedziak et al., 2014). Alternatively, a recent study posited that FtsZ simply serves as a scaffold and that the constrictive force on the membrane is provided by the inward growing cell wall (Coltharp et al., 2016). This model was suggested by the observation that the rate of inward cell wall growth is limited by the rate of cell wall synthesis but not by the GTP hydrolysis rate of FtsZ.

In order to explore these different conceptual models, we modified our coarse-grained simulation software for the Gram-negative bacterial cell wall, REMODELER 1, to create REMODELER 2, which allowed us to test different mechanistic hypotheses of how inward cell wall growth might occur during division. We found that simply restricting the enzyme complexes to the midcell resulted in elongation without constriction, even with a make-before-break mechanism of PG remodeling, suggesting that cell wall growth alone is not sufficient to drive Gram-negative bacterial cell division. We found that a constrictive force at the midcell did result in cell wall division when the force was sufficiently large to initially constrict the midcell past the diameter of the unpressurized sacculus. If the constrictive force was slightly less, sufficient to constrict the midcell sacculus into a relaxed state, the addition of a make-before-break mechanism was now effective in facilitating division. These results are summarized in Movie S1. Due to the difficulty of describing dynamic 3D processes in words and static images, we recommend readers watch the movie in full before proceeding.

## Results

### Cell wall synthesis at the midcell

To adapt REMODELER 1 (Nguyen et al., 2015) into REMODELER 2 to study cell division, we added several features: (1) PG synthesis complexes were organized at the midcell, (2) a constrictive force could be implemented, and (3) the enzymes could build a multi-layered cell wall.

To simulate PG insertion during cell division, we built a starting PG sacculus and initiated four PG synthesis enzyme complexes randomly around the circumference within 10 nm of the midcell. To reduce the computational cost, we simulated a short cylindrical section (a midcell) of a miniaturized cell wall. Specifically, the starting PG cylinder was composed of40 glycan hoops with each hoop consisting of 400 beads, representing 400 tetrasaccharides (Fig. 1A). The average strand length was set to be 4b tetrasaccharides, which is within the range of 11–16 tetrasaccharides reported experimentally (Glauner et al., 1988; Harz et al., 1990). As the distance between adjacent tetrasaccharides was *L*_*g*_ = 2 nm (Nguyen et al., 2015), the unpressurized PG cylinder had a radius of 127.5 nm (Fig. 1A). Under a turgor pressure *p*_*tg*_= 3 atm, the cylinder expanded to a radius of 137.5 nm (Fig. 1B). To minimize any potential effects of changing the glycan strand length on the sacculus radius, new glycan strands were also constrained to 4b tetrasaccharides on average when they were incorporated into the existing PG network. As in our previous simulations of PG remodeling (Nguyen et al., 2015), here we also assumed that the PG synthesis enzyme complexes insert new glycan strands in pairs (Fig. S1) and that the enzyme complexes can only act on PG substrates in their close vicinities, and cannot stretch the new pair of strands or pull them forward to crosslink them to distal peptides. At this stage of our model, we tested the hypothesis that simply organizing the PG synthesis complexes at the midcell can cause constriction (Meier and Goley, 2014; Eun et al., 2015). In our simulations, however, insertion of new PG only elongated the cylinder without changing its radius (Fig. 1C-D), suggesting that additional factors are needed to induce division.

**Figure 1.**
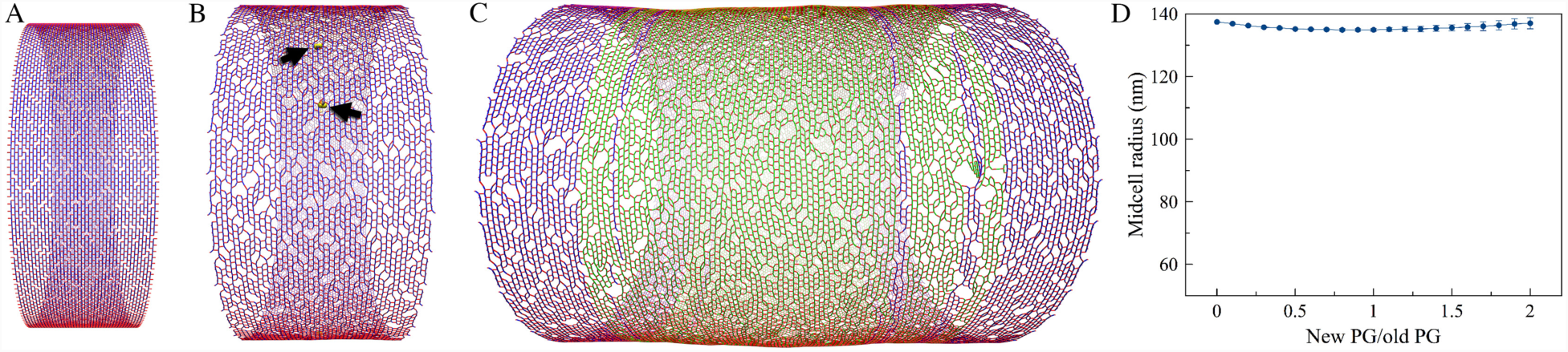
Cell wall synthesis at the midcell. Existing glycan strands are visualized in blue, new strands in green, peptide crosslinks in red. The same color scheme is used for all other figures containing simulation snapshots. (A) A snapshot of the starting PG cylinder in a relaxed state. The cylinder was composed of 40 hoops of glycan strands with 400 tetrasaccharides per hoop and had a relaxed radius of 127.5 nm. (B) A snapshot of the pressurized PG cylinder which was expanded to a radius of 137.5 nm under a turgor pressure of 3 atm. The arrows indicate PG synthesis enzyme complexes placed randomly at the midcell. (C) A snapshot of the PG cylinder after elongation to three times its original length by insertion of new PG. (D) The profile of the midcell radius with respect to the amount of new PG inserted, showing that PG synthesis at the midcell alone only elongated the cylinder without constricting it. The average of 4 simulations is shown. Error bars indicate standard deviation.

### PG remodeling under a constrictive force

Since FtsZ filaments have been proposed to exert a constrictive force on the membrane, we implemented a constrictive force at the midcell to see if this allowed new PG to be incorporated in smaller hoops at the constriction site (see **Methods/Constriction force**). Initially, the constrictive force made the midcell smaller before new PG was inserted (Fig. 2A). As new PG was inserted, further reduction of the midcell radius did not occur if *F*_*c*_, the constrictive force divided by the sacculus circumference, was smaller than 20 pN/nm (Fig. 2B, 2D). The midcell did continue to reduce in size if *F*_*c*_ was larger than 20 pN/nm (Fig. 2C, 2D). We found that at the transition point *F*_*c*_ ~ 20 pN/nm, the force initially constricted the midcell into a relaxed state where its radius was equivalent to that of an unpressurized cell, 127.5 nm (Fig. 2E). Therefore, a constrictive force alone can drive division if it is sufficiently large to initially bring the midcell into a compressed state, i.e. reduce the midcell radius to less than that of a relaxed sacculus. Note also that our findings were limited to *F*_*c*_ less than ~32 pN/nm since a larger force buckled the cell wall, making the simulation unstable.

**Figure 2.**
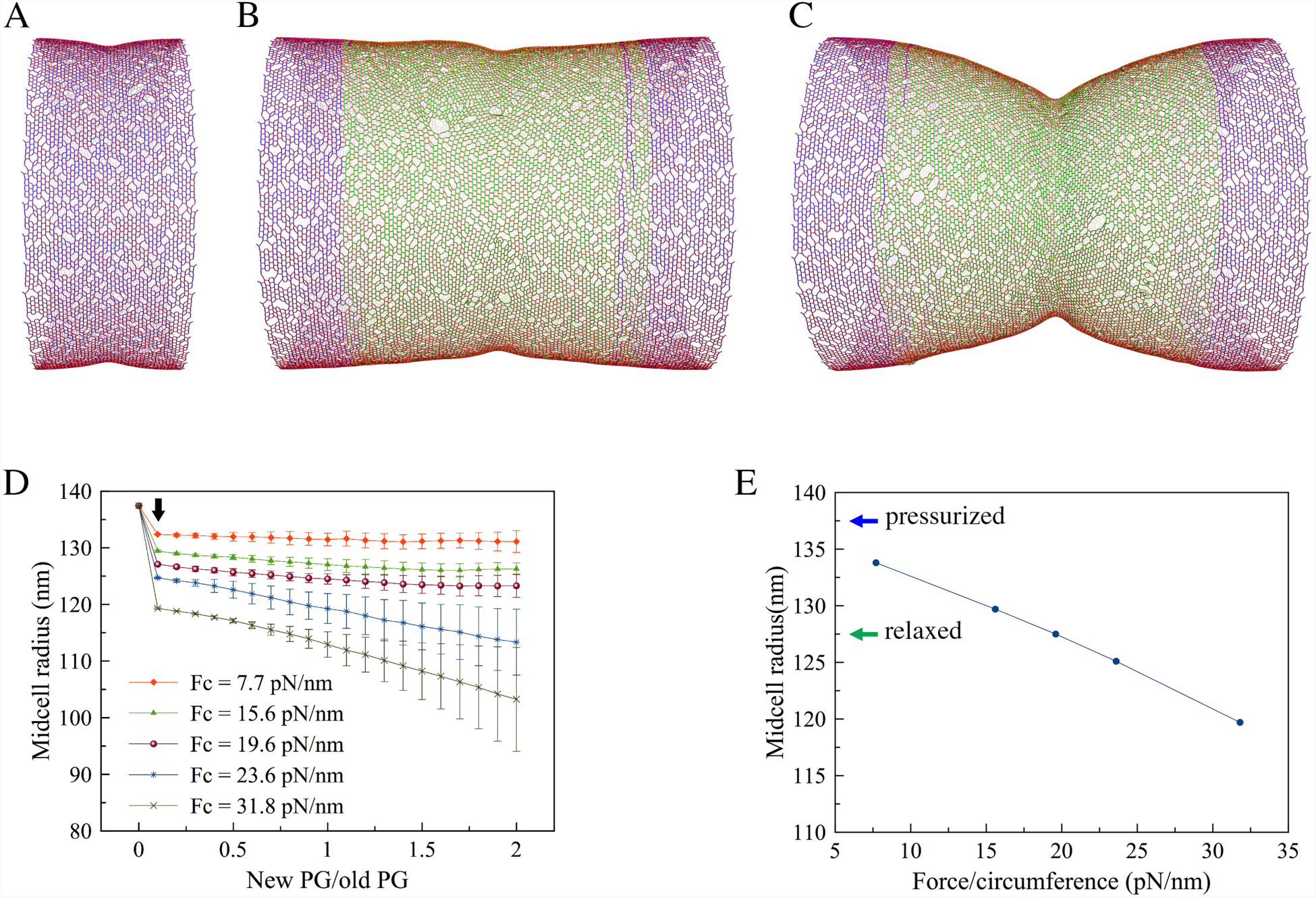
Cell wall synthesis under constrictive force. (A) A snapshot of the PG cylinder showing that the force caused an initial constriction at the midcell before insertion of new PG. (B) A snapshot of the sacculus after new PG was inserted showing that when the force per circumference was *F*_*c*_= 15.6 pN/nm, further constriction did not occur as new PG was inserted. (C) When *F*_*c*_ = 31.8 pN/nm, constriction occurred as new PG was inserted. (D) Profiles of the midcell radius with respect to the amount of new PG inserted show that PG insertion-induced constriction occurred only if *F*_*c*_ was larger than 19.6 pN/nm. The arrow indicates the initial constriction by the force before new PG insertion started. Each trace presents the average of 4 simulations. Error bars indicate standard deviation. (E) The dependence of the midcell radius on the force before new PG was inserted shows that the midcell became relaxed at *F*_*c*_ = 19.6 pN/nm. The blue arrow indicates the radius before the force was applied. The green arrow indicates the radius of a relaxed cell wall.

We next analyzed in detail how a constrictive force might drive cell wall division. Without the constrictive force, the cell wall radius was maintained as new glycan beads were perfectly matched one-to-one with the existing template (Fig. 3A). Applying a small constrictive force (*F*_*c*_ < 20 pN/nm) squeezed the midcell and pulled the enzymes and the two new strand tips closer to the default template crosslink, but this did not interrupt the one-to-one template matching between the new beads and the existing beads and therefore did not reduce the midcell radius (Fig. 3B). On the other hand, in the presence of a large force (*F*_*c*_ > 20 pN/nm), the enzymes were pulled past the default template crosslink, skipping it and crosslinking the two new beads to a new template that was upstream of the skipped template (Fig. 3C). Due to these skipping events, the two new PG hoops had fewer PG beads than the existing hoops, making the midcell radius smaller.

**Figure 3.**
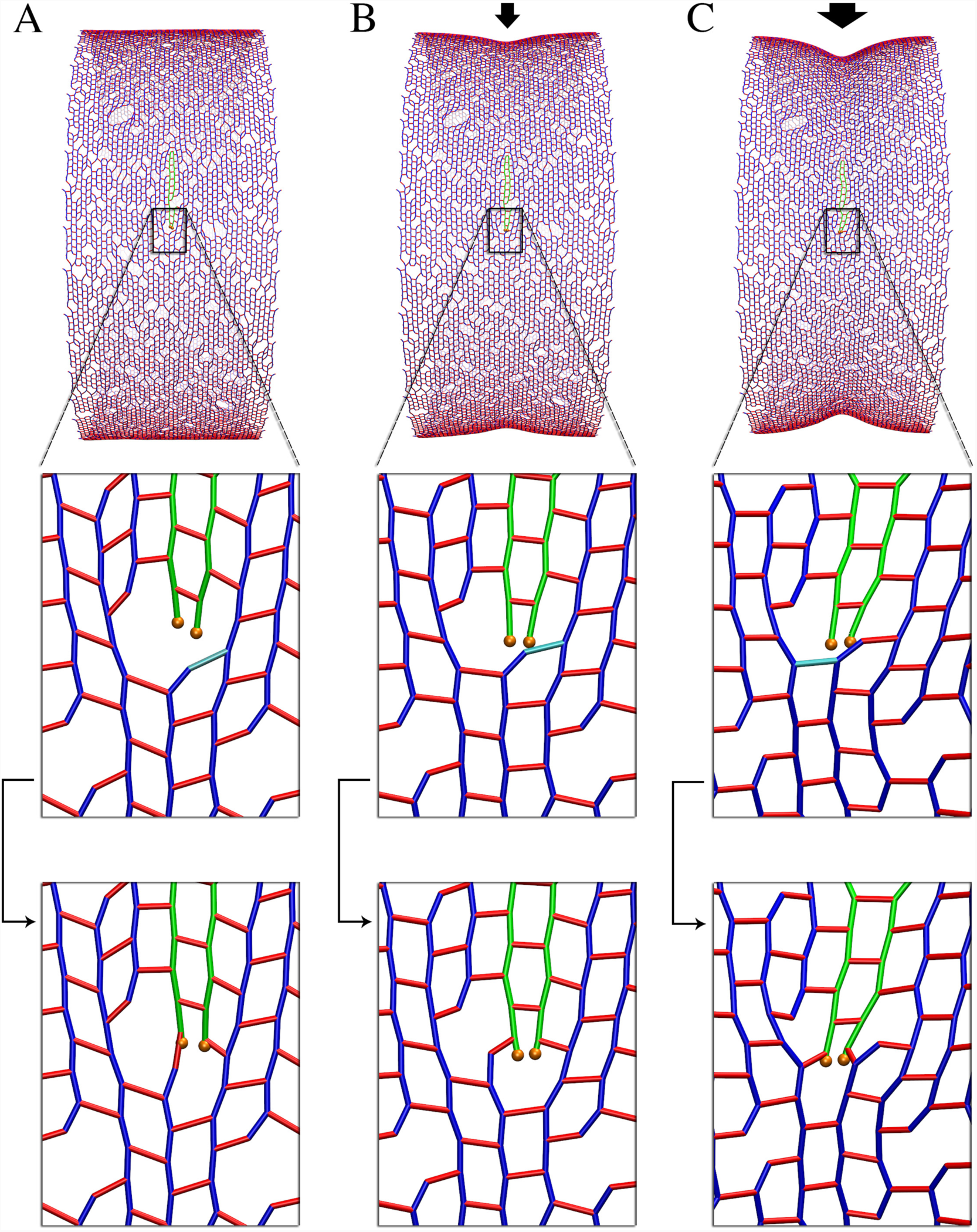
Detailed effect of the constrictive force. The template crosslink is highlighted in cyan and the arrows between the zoomed-in views in the bottom two rows indicates the time sequence of events. (A) Without the force, new PG was matched one-to-one with the existing template. (B) In the presence of a small force, the midcell was squeezed, pulling the new strand tips closer to the template crosslink, but the new PG was still matched with the default template. (C) In the presence of a large force, the new strand tips were pulled past the default template, therefore skipping it, and an upstream crosslink became the new template. Note that several such template-skipping events occurred along each complete hoop of new PG.

**Figure 4.**
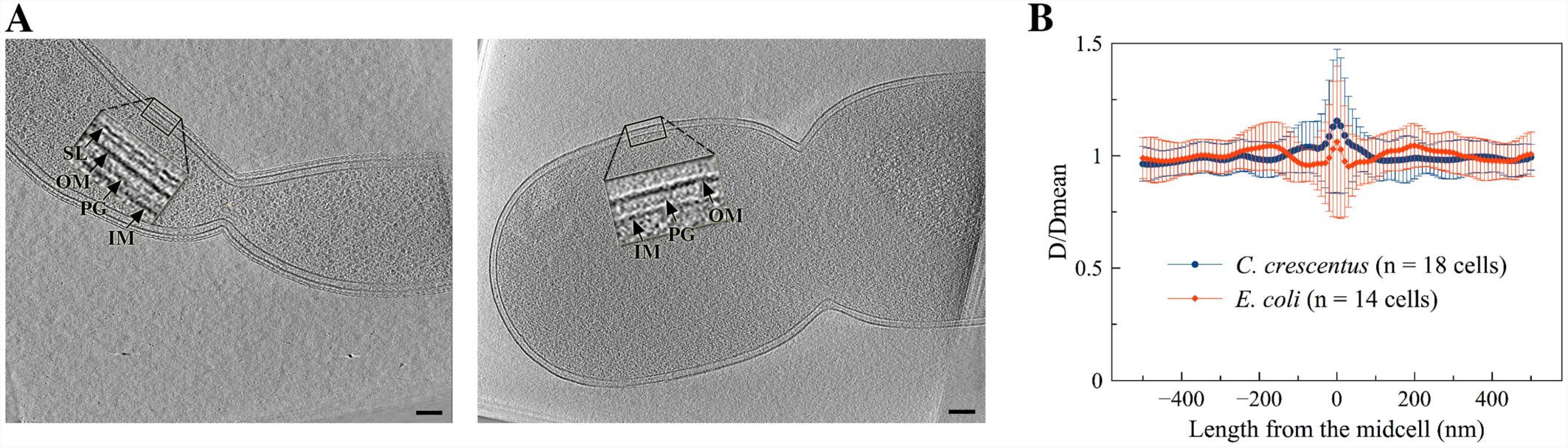
Electron cryotomography of dividing Gram-negative bacterial cells. (A) Representative tomographic slices through dividing *Caulobacter crescentus* (left) and *Escherichia coli* cells (right). Arrows point to the S-layer (SL), outer membrane (OM), peptidoglycan (PG), and inner membrane (IM). Scale bars indicate 100 nm. (B) Calculation of the distance *D* between the two membranes shows that it is maintained throughout the invagination site at the midcell. Error bars indicate standard deviation.

### PG remodeling under a make-before-break mechanism

Next, we explored if and how cell wall growth alone could be sufficient to drive cell division without the presence of a constriction force. Conceptually, this can occur with a make-before-break mechanism, in which the cell wall synthesis machinery adds one or several new PG layers that form a temporary septum underneath the existing PG layer (make) before hydrolases cleave the constraining peptide crosslinks above these new PG layers (break). If many layers are built in before any bond on the surface is hydrolyzed, the hoops of PG in the inner layer can potentially be made from fewer PG beads. While it is clear that Gram-positive bacteria divide by making a thick septum across the width of the cell, it has only recently been speculated that Gram-negative bacteria might also adopt this septation scheme, but with a thinner septum (Erickson, 2017). (For clarity, we use septum here to refer to the new PG layers beneath, but not including, the existing layer.)

To determine if such a septum exists in dividing Gram-negative bacterial cells, we examined 3D electron cryotomograms of intact frozen-hydrated cells of six speciesa *Caulobacter crescentus, Escherichia coli, Proteus mirabilis, Myxococcus xanthus, Cupriavidus necator*, and *Shewanella oneidensis*. In all cases, we could not discern any thickening of the wall at the dividing midcell that might indicate the existence of a thin septum (Fig. 4A; Fig. S2). We also observed that the distance between the inner and outer membranes remained constant throughout the midcell (Fig. 4B). We therefore concluded that if a thin septum exists at the dividing midcell, it must be thinner than ~4 nm, the resolution of the electron cryotomograms (Gan and Jensen, 2012). Accordingly, in our simulations, we limited septum thickness to one layer of PG.

We simulated a make-before-break mechanism by decoupling PG synthesis (transglycosylation and transpeptidation) from PG hydrolysis (endopeptidation). Specifically, cleavage of existing peptide crosslinks was blocked until complete hoops of new glycan strands were crosslinked into the PG network underneath these crosslinks (Fig. 5A-D). Note that how this might occur at a molecular level remains unclear. The rate of endopeptidases was controlled so that only ~one layer of new PG was present underneath the existing layer (see **Methods/Make-before-break mechanism**). To mimic the volume exclusion effect between the outer and inner layers, before existing peptide crosslinks were cleaved, a repulsive force between these crosslinks and the new glycan strands was applied to separate them to a distance of 2 nm, the estimated thickness of one PG layer. To study the effect of the make-before-break mechanism alone, we did not apply a constrictive force. Simulation results showed, however, that this make-before-break mechanism did not reduce the midcell radius (Fig. 5E, 5F). We found that once the existing crosslinks above the new PG hoops were cut, the inner hoops expanded to the size of the existing hoops (Fig. 5C, 5D), indicating that new hoops were made of a similar number of beads as existing hoops.

**Figure 5.**
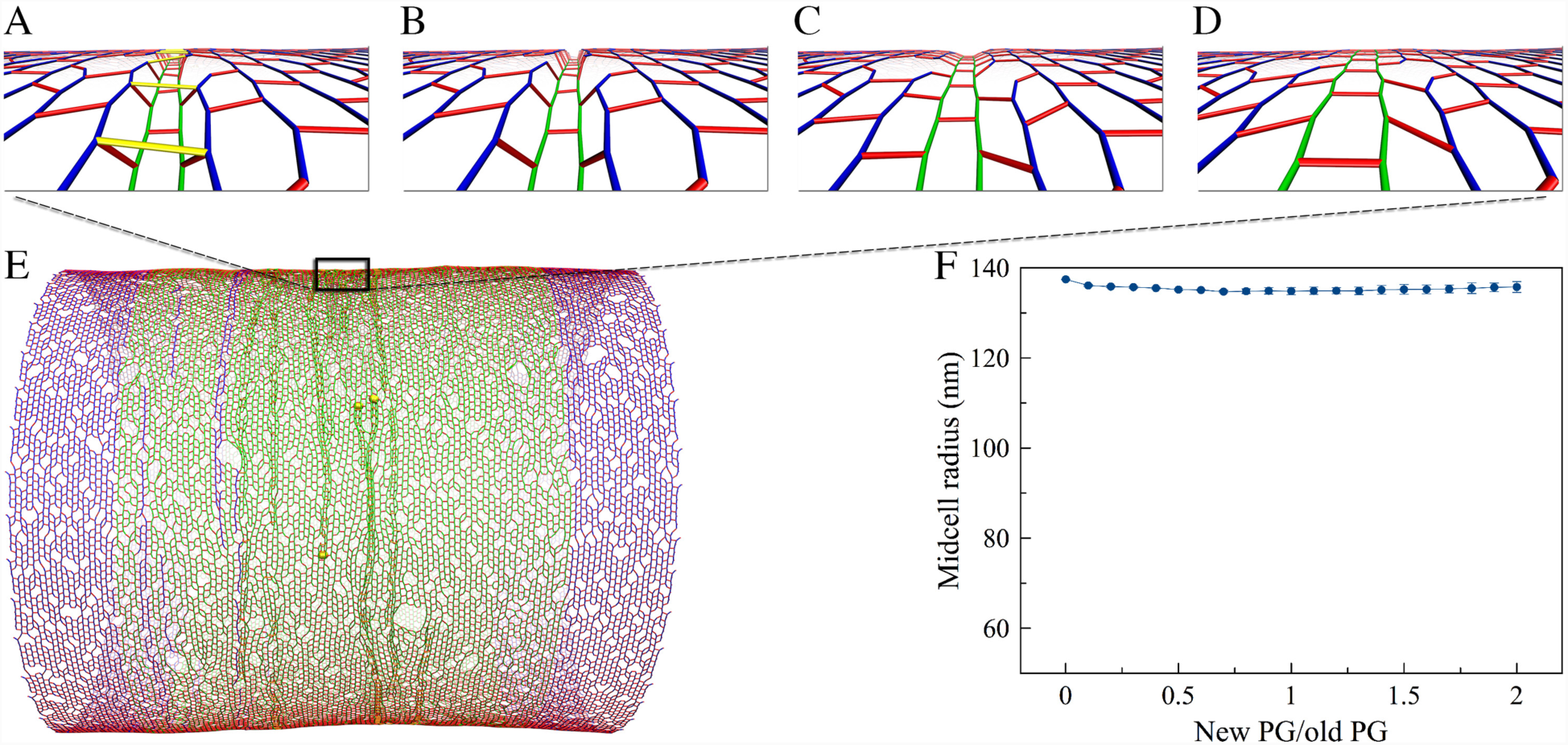
Cell wall synthesis with a make-before-break mechanism. (A) – (D) show the make-before-break mechanism occurring in sequencea (A) new cell wall was made underneath the existing network. The peptide crosslinks above the new PG are highlighted in yellow. (B) The highlighted crosslinks were cleaved after complete hoops of new PG were made. (C) and (D) show the relaxation of the network after the cleavage event. (E) A snapshot of the PG cylinder shows that constriction did not occur with insertion of new PG. (F) The profile of the midcell radius with respect to the amount of new PG inserted. The averages of 4 simulations is shown. Error bars indicate standard deviation.

We then analyzed why this make-before-break model failed. Conceptually, if turgor pressure is not present, all the PG beads are evenly spaced at *L*_*g*_ = 2 nm (the length of one tetrasaccharide) on the same hoop. Matching the beads on the smaller hoop to those on the larger hoop would then create mismatches (Fig. S3). With a difference in radius Δ*r* = 2 nm, the new hoop’s circumference would be 2*π*Δ*r* = 12.56 nm shorter than that of the existing hoop. Since there are *N*_*b*_ = 400 beads on the existing hoop, the average mismatch per bead would be Δ*s* = 2*π*Δ*r/N*_*b*_ ~0.0314 nm, which is small compared to the distance between the adjacent beads. However, if the first pair of beads on the two new strands (as discussed above, we assumed new strands are synthesized in pairs) are in register with their templates on the existing hoops (Fig. S3B), the second pair of new beads would be positioned ahead of their template by a distance Δ*s* = 0.031b nm. After ~ 30 pairs of beads are added to the new strands, the accumulated shifting of the strand tips would become 30Δ*s* ~1 nm, about half the distance between adjacent beads (Fig. S3C). At this point reaching backward for the default crosslink template would become unfavorable, so instead the enzymes might skip the default template and reach forward for an upstream crosslink on the adjacent track (Fig. S3D). If such a skip occurred, the new strand tips would now trail their template by a distance of ~1 nm. It would then take another 30 beads for the new tips to catch up and once again be in register with their template (Fig. S3E). The cycle would then continue, leading to template skipping every 60 beads and a complete new hoop of only 394 beads, 6 beads less than the existing hoops. This would decrease the diameter of the midcell.

In the presence of turgor pressure, however, the cell wall expanded as peptide crosslinks were stretched and tilted away from the long axis of the cylinder (Fig. 6). As a result, beads were no longer evenly spaced on the same hoop. At breaks between glycan strands in the hoop, the gap between the adjacent peptide crosslinks expanded from 2*L*_*g*_ = 4 nm (Fig. 6A) to ~ 6.2 nm (Fig. 6B) as terminal peptides tilted an average of 30° (SD = 15°) (Fig. 6C). We observed that right before encountering a glycan break on the existing strands, which occurred every ~14 beads, the new strand tips had gotten ahead of their templates by an accumulated distance *s*_*a*_ = 14Δ*s* = 0.44 nm (Fig. 6D). At the glycan break, though, this small progress was more than offset by the 2.2-nm turgor pressure-induced expansion of the gap (Fig. 6E). At this stage, the new strand tips even fell behind their templates (Fig. 6F). This lag did not accumulate, however, because new strands also terminated, at which point the next new strands were pulled forward (Fig. 6G). This meant that template beads were not skipped, the new hoops had the same number of beads as the existing hoops, and constriction did not occur.

**Figure 6.**
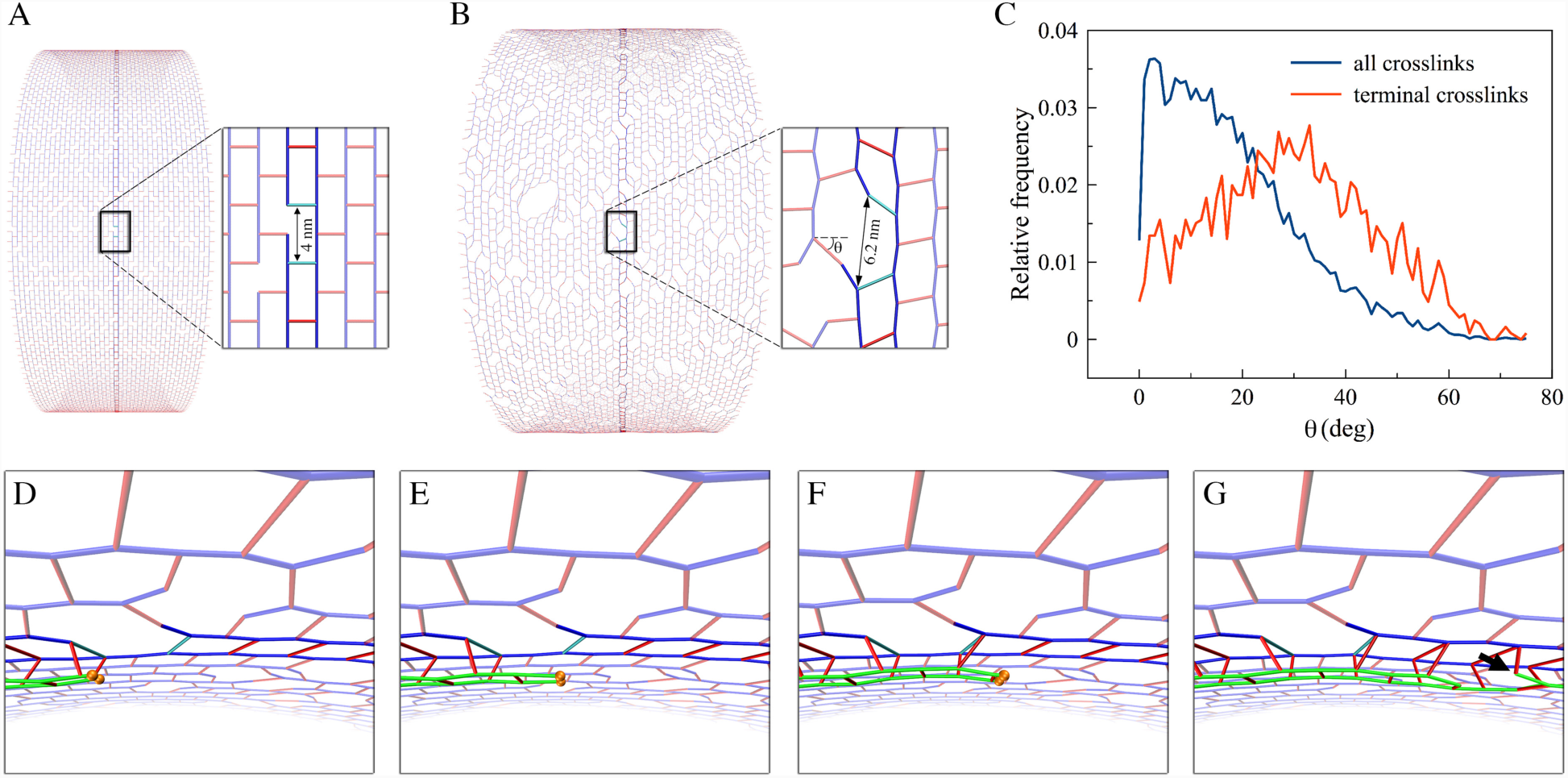
Effect of turgor pressure on the make-before-break mechanism. Two adjacent peptide crosslinks at a glycan break in a hoop are highlighted in cyan for visualization. (A) Without turgor pressure, the PG was well-ordered and the distance between the highlighted crosslinks was 4 nm. (B) Under turgor pressure, the cylinder expanded, peptides tilted, and the distance between the highlighted crosslinks increased to ~6.2 nm. *θ* depicts the tilting angle of a peptide crosslink. (C) Histogram of the tilting angles of all the peptide crosslinks (blue) and those connecting glycan termini (red). (D) An oblique view showing insertion of two new glycan strands in a make-before-break mechanism. When the new strand tips reach the first highlighted crosslink, they get ~0.4 nm ahead of their templates. (E) At the second highlighted crosslink, this small gain is offset by the additional 2.2 nm enlargement of the distance between the two crosslinks. (F) After the second highlighted crosslink, the strand tips fall behind their templates. (G) A break in the new glycan strands (indicated by the arrow) pulls the new strands forward, preventing them from falling behind their templates.

Since cell expansion due to turgor pressure varies with cell size (Fig. S4A), the difference in radius of the cell wall Δ*r* between a relaxed cell and a pressurized cell can be negligible for cells of sufficiently small sizes. For example, for cells with circumference of fewer than 150 tetrasaccharides (corresponding to a diameter of ~100 nm), Δr would be smaller than 2 nm, which is the assumed thickness of one PG layer (Fig. S4B). In this case, a make-before-break mechanism could plausibly drive division in the absence of a constrictive force. In addition to cell width, turgor pressure may vary between cells. While most studies report turgor pressure in the range of 2–4 atm in Gram-negative bacteria (Reed and Walsby, 1985; Koch and Pinette, 1987; Cayley et al., 2000), a turgor pressure as low as 0.3 atm has been reported (Deng et al., 2011). At this low turgor pressure, cells of circumference smaller than 300 tetrasaccharides would expand less than 2 nm in radius (Fig. S4B). Still, for a make-before-break mechanism to be effective in a cell the size of an *E. coli* (~1500 tetrasaccharides in circumference), the turgor pressure would need to be on the order of 0.03 atm to make the radius expansion negligible (Fig. S4C).

### Make-before-break in the presence of a constrictive force

Our observations suggested that in order for the make-before-break PG remodeling mechanism to be effective in constriction, the midcell must be in a relaxed state. We reasoned that a constrictive force squeezing the midcell could create this condition by restoring the gaps between adjacent peptide crosslinks at glycan breaks from ~6.2 nm to their relaxed size of ~4 nm (Fig. 7A). Implementing a constrictive force per circumference length *F*_*c*_ smaller than ~20 pN/nm resulted in an initial constriction that did not completely relax the midcell. Predictably, in this scenario, insertion of new PG with a make-before-break mechanism did not cause further constriction (Fig. 7C). When *F*_*c*_ was ≳ 20 pN/nm, the midcell was completely relaxed or even squeezed to a smaller radius than the unpressurized cell (Fig.2E). As we expected, in this condition make-before-break PG remodeling could now reduce the midcell radius (Fig. 7B, 7C). Note that while the constriction force alone could drive division if the magnitude of *F*_*c*_ was larger than 20 pN/nm, reducing the midcell to a compressed state (Fig. 2E), in the presence of the make-before-break mechanism, division started to occur at *F*_*c*_ = 20 pN/nm since reducing the midcell to a relaxed state was sufficient.

**Figure 7.**
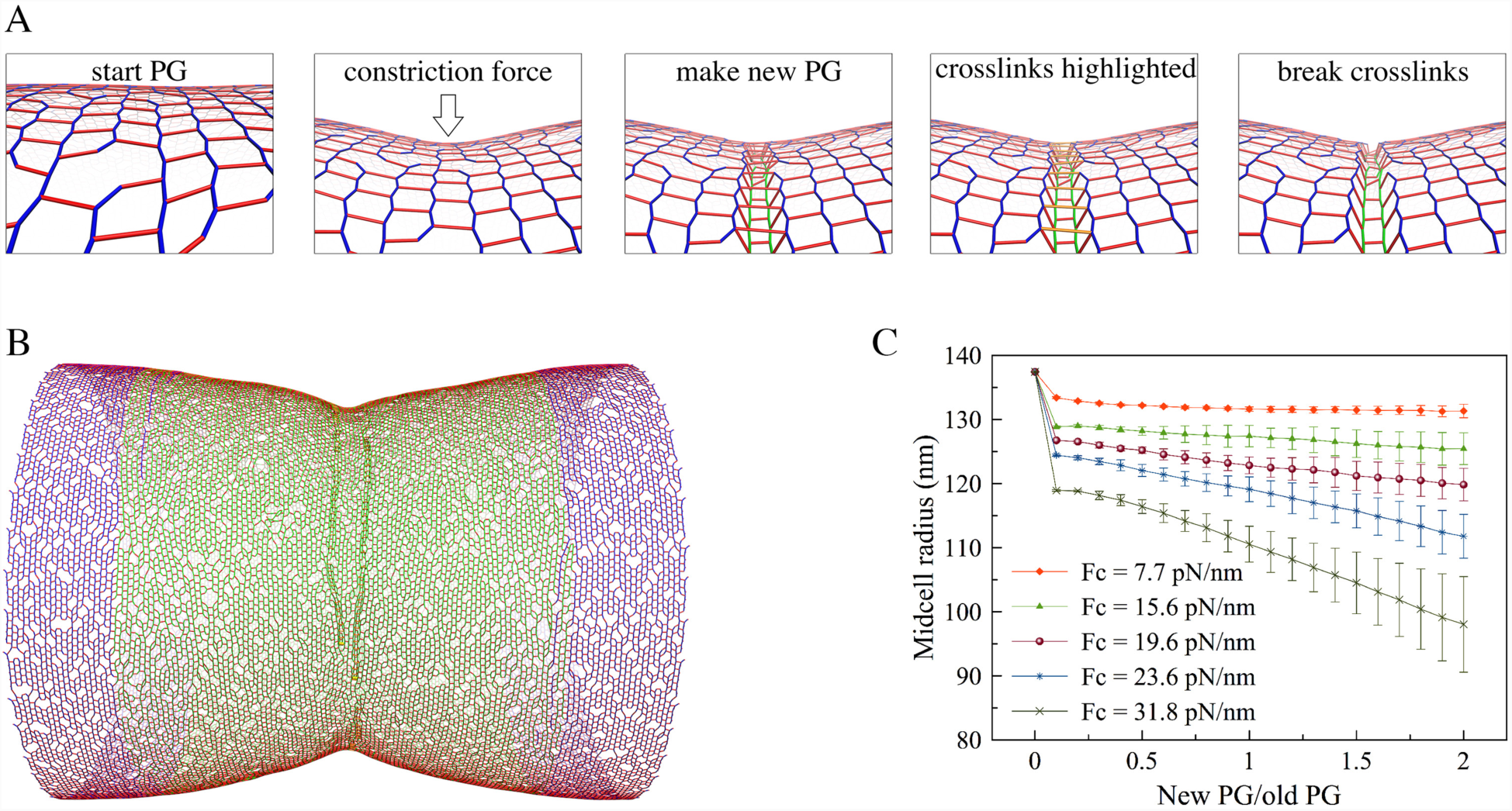
Cell wall synthesis in the presence of both a constrictive force and the make-before-break mechanism. (A) Representative snapshots of a simulation sequencea the constriction force caused an initial constriction, and new PG was made underneath the existing network in complete hoops before crosslinks (highlighted in orange) above the new hoops were cleaved. (B) A snapshot of the sacculus showing that constriction occurred upon insertion of new PG when the force per circumference *F*_*c*_ = 23.6 pN/nm. (C) Profiles of the midcell radius with respect to the inserted amount of new PG. Each trace shows the average of 4 simulations. Error bars indicate standard deviation.

## Discussion

Together, our results demonstrate how 3D modeling of molecular details can provide insights into complex symmetry-breaking processes such as cell division. By simulating how rod-shaped Gram-negative bacteria could divide their cell walls, we found that a constrictive force is the key factor driving constriction, while cell wall remodeling by a make-before-break mechanism can only facilitate the process. We note, however, that our results are limited by three assumptions: (1) PG synthesis enzymes only act on substrates locally in a complex, (2) a force generator (presumably FtsZ) provides a constrictive force at the midcell during cell division, and (3) the cell can remodel its cell wall by a make-before-break mechanism in which new hoops of PG are made inside the existing hoops before peptide crosslinks on the old hoops are cleaved. Note that the original make-before-break model was proposed for individual glycan strands, not complete hoops of strands (Koch, 1990; Höltje, 1993). While experiments are needed to validate or refute these assumptions, our work provides, to our knowledge, the first *in silico* insights into how cells might employ different driving forces to divide their cell walls.

### Conceptual models of cell wall division

To decrease the radius of the midcell, the cell needs to add smaller and smaller hoops of new PG. Conceptually, this can occur by three models. (1) Because existing PG hoops are stretched by turgor pressure, if by some unknown mechanism new PG hoops are stretched even further at the time they are incorporated between existing hoops, the new hoops would have fewer PG beads and therefore become smaller as the system relaxes. (2) If a mechanism exists to initially compress existing hoops at the midcell and new, relaxed PG hoops are incorporated between these existing hoops, the new hoops would have fewer beads and become smaller upon relaxation. (3) If a mechanism exists to initially relax existing hoops at the midcell and new, also relaxed, PG hoops are made inside existing PG hoops, the new hoops would again have fewer beads. Here we showed that Model 2 is plausible with a constriction force alone and Model 3 is plausible with a combination of a constriction force and a make-before-break mechanism. We did not simulate Model 1 as we judge it unlikely to occur in real cells. Nevertheless, we cannot currently rule out the possibility that an unknown force pulls the enzymes forward, stretching the new glycan strands before incorporating them. Nor can we rule out the possibility that cells might divide by a completely different mechanism that has not yet been discussed in the literature.

### The role of a constrictive force

FtsZ filaments have been shown to be able to constrict liposomes (Osawa and Erickson, 2013), but it is unclear if they exert a constrictive force on the membrane *in vivo* and if this force is required for cell division. Here we found *in silico* that to make new PG hoops smaller than existing hoops, the midcell needs to be initially constricted to at least a relaxed state, with further constriction occurring only after new PG is inserted. In this model, the constriction rate is limited by the slower of either the force generator (presumably FtsZ) or the PG synthesis rate. Therefore, the finding by Coltharp et al. that the inward growth rate of the cell wall is limited by the rate of PG synthesis but did not change even when the GTP hydrolysis rate of FtsZ was reduced 90% (Coltharp et al., 2016) might simply reflect that the PG synthesis rate is much slower than the action of FtsZ. Indeed, it has been reported that an FtsZ mutant with a GTP hydrolysis rate 3% that of wild type FtsZ resulted in very slow growth of colonies (Redick et al., 2005).

In our model, the total initial constriction force needed to be at least ~15 nN (corresponding to a force per circumference *F*_*c*_ ~20 pN/nm) to enable cell wall division. Assuming that the constrictive force is generated by FtsZ, each monomer of which has been estimated by molecular dynamics simulations to generate 30 pN (Hsin et al., 2012), our estimated force is equivalent to the action of 500 FtsZ monomers, which could form a continuous filament ~2.2 μm long or 15 filaments of an average length of 150 nm. This is reasonable, considering an estimated ~5-7,000 FtsZ molecules per cell measured in *E. coli* (Erickson et al., 2010) and the fact that our simulated sacculus is a third the size of an *E. coli* cell. Note that it has recently been speculated that excess membrane synthesis might also generate a constrictive force (Osawa and Erickson, 2018).

### Can cell wall growth alone drive constriction?

While cell wall growth has been speculated to partially or primarily drive constriction during cell division (Meier and Goley, 2014; Coltharp et al., 2016), our simulations showed that cell wall growth via a make-before-break mechanism failed to cause cell division in the absence of a constriction force. We found that for the make-before-break mechanism to have an effect, the new PG hoops must be made inside relaxed existing hoops and therefore a constriction force is needed to initially relax the midcell. In theory, the need for a constriction force could be bypassed if the enzymes could make a multi-layered septum. Once the septum thickness was equal to or larger than the difference in radius between the pressurized cell and the relaxed cell, the innermost layer of the septum would be in a relaxed state, allowing the next PG layer to be made of hoops containing fewer PG beads (Fig. S5). For this scenario to occur, in the case of our modeled cell wall, which had a radius of 127.5 nm when relaxed and 137.5 nm when pressurized, the septum would have to contain at least five PG layers (assuming each layer is 2 nm thick). By this logic, a cell the size of an *E. coli*, whose circumference is ~1500 tetrasaccharides, would need a septum ~65 nm thick for this mechanism to drive division (Fig. S4A-B). However, we saw no such septa in our electron cryotomograms of dividing cells. We therefore think it unlikely that this mechanism is the primary driver of cell division.

## Methods

### Simulation of cell wall synthesis

Here we only briefly describe our simulation system. For a more detailed description, please see our previous paper (Nguyen et al., 2015).

### Cell wall

We coarse-grained the cell wall such that each glycan strand is represented as a chain of beads, each bead represents one tetrasaccharide, and the peptides attached to the beads alternate between the left and right sides. Adjacent glycan beads are connected by springs of a relaxed length *l*_*g*_ = 2 nm and a spring constant *k*_*g*_ = 5.57 nN/nm. The bending stiffness of the strand is *k*_*b*_ = 8.36 · 10^-20^ J and the relaxed angle at the beads is *θ*_0_ = 3.14 rad. We modeled peptide crosslinks as worm-like chains such that if the peptide end-to-end extension *x* is larger than *x*_*0*_ = 1.0 nm the following force is applied

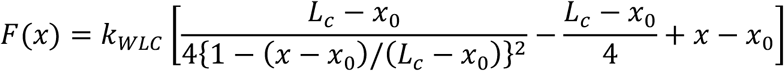

where *L*_*c*_ = 4.8 nm is the contour length of the peptide crosslink and *k*_*WLC*_ = 15.0 pN/nm is the force constant.

Previously, in order to reduce the computational cost, most of our simulations started with a small sacculus with a circumference composed of 100 tetrasaccharides (Nguyen et al., 2015). In the current simulations, to allow the midcell radius to constrict over time, we used a starting sacculus with a circumference of 400 tetrasaccharides. To reduce the computational cost, since PG remodeling only occurs at the midcell during cell division, we removed the two caps of the starting sacculus and built a cylinder only 40 glycan hoops wide.

### PG remodeling enzymes

Four enzyme complexes were added at the midcell. In each complex, three types of PG remodeling enzymes are explicitly represented as beads and a house-keeping enzyme that cleaves the long tails of glycan strands is implicitly implemented. Specifically, there are two transglycosylases that each synthesizes a glycan strand (so two strands emerge from the complex) (Fig. S1). On average, each transglycosylase adds a tetrasaccharide bead every 103 time steps. Transglycosylase then translocates to the strand tip to be ready to add another bead. Note that previously we hypothesized that transpeptidation facilitates translocation. Specifically, the probability of translocation is once every 10^6^ time steps if the last-added bead is not crosslinked, but this probability becomes once every 3 10^4^ time steps after the last-added bead is crosslinked (these numbers were arbitrarily chosen because we were not aware of experimentally-reported enzyme rates) (Nguyen et al., 2015). Considering that the modeled sacculi in our current simulations were 4 times larger than those in our previous simulations, to speed up the current simulations, we increased the probability of transpeptidation-facilitated translocation 10 times to become once every 3 103 time steps. To maintain an average glycan strand length of 14 tetrasaccharides, the termination probability of strand elongation is also increased two-fold to once every 2 10^6^ time steps. Note also that in our previous simulations, interactions between transglycosylases and outer-membrane lipoproteins LpoA and LpoB were implemented that prevented the transglycosylase-lipoprotein complex from crossing through glycan strands or peptide crosslinks. To enable the make-before-break mechanism, these transglycosylase-lipoprotein interactions were removed from the current model, allowing transglycosylases to freely move across strands and crosslinks.

One endopeptidase exists in each enzyme complex to cleave existing peptide crosslinks. In our previous simulations, when an endopeptidase diffused across a crosslink, the enzyme cleaved the crosslink with a probability of 0.1. To speed up our current simulations, every 10 time steps, if the distance from the endopeptidase to a crosslink is within 3.0 nm, the enzyme captures then cleaves the crosslink. If there are multiple crosslinks within this reaction distance, the probability of crosslink i being chosen is calculated as

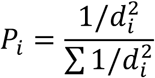

where *d*_*i*_ is the distance from the endopeptidase to crosslink *i*.

There are three transpeptidases in each complex, one crosslinking the two new strands to one another and the other two crosslinking the pair to the existing network (Fig. S1). Previously, the probability of a transpeptidase capturing a peptide of a PG bead at a distance dwas given as *P*_*tp*_ = (1 - *d*/*d*_0_)^2^ where *d*_0_ = 2.0 nm was the reaction distance. To speed up the current simulations, *d*_0_ is increased to 3.0 nm. Note that increasing the modeled rates of the enzymes did not change the principles driving cell wall remodeling in our simulations.

### Turgor pressure

As in our previous simulations, turgor pressure was chosen to be *P*_*tg*_ = 3.0 atm. The force on the pressurized sacculus of volume *V* is calculated as *F*_*tg*_ = -∇*E*_*vol*_ where *E*_*vol*_= −*P*_tg_*V* is the work done by turgor pressure to inflate the sacculus.

### Constriction force

When a constriction force is applied at the midcell, it creates an inward pressure *P*_*c*_ satisfying *F*_*c*_

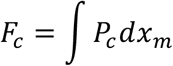

where *F*_*c*_ is the constriction force divided by the circumference length and *x*_*m*_ is the distance from the midcell. To model the force, if the absolute value of *x*_*m*_ is less than 50 nm, a constriction pressure [inline]is applied, where *σ*= 10 nm and *P*_0_ is calculated using the following equation

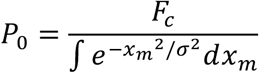

### Make-before-break mechanism

To implement the make-before-break mechanism, existing peptide crosslinks are marked as gconstraining crosslinksg once new glycan strands are formed underneath the crosslinks. Cleavage of a “constraining crosslink” is delayed for 10^6^ time steps. After that, cleavage can occur by four scenariosa (a) an endopeptidase is within 2 nm of the constraining crosslink, (b) two PG layers exist beneath the crosslink, (c) a random cleavage with a probability of once every 10^7^ time steps, or (d) the constraining crosslink has existed for 10^9^ time steps. We found that these probabilities resulted in a septum ~1 PG layer thick.

In real cells, different PG layers would be separated due to the volume exclusion effect. To mimic this effect in our model, if a glycan strand is underneath a constraining crosslink and their separation d is less than the thickness of one PG layer *t*_*g*_ = 2.0 nm, a repulsive force *F*_*r*_ = *k*_*r*_(*t*_*g*_ – *d*) is exerted on the crosslink and the two PG beads that are closest to the crosslink, where the force constant *k*_*r*_ was arbitrarily chosen to be 200 pN/nm.

### Electron cryotomography

Bacterial strains were grown and imaged as described: *Caulobacter crescentus* (Yao et al., 2017); *Escherichia coli* (Pilhofer et al., 2011); *Myxococcus xanthus* (Chang et al., 2016); *Cupriavidus necator* (Beeby et al., 2012); *Shewanella oneidensis* (Kaplan et al., 2018).

### Measurement of the distance between the inner and outer membranes

A tomographic slice 10 nm thick through a central plane along the long axis of the cell was captured using the IMOD software (Kremer et al., 1996). Each membrane (inner and outer) was manually traced and represented by a set of points evenly spaced along the line. This process was repeated for the membranes on the opposite side of the cell. The location of the midcell was determined by the shortest distance between the two traces of the inner membrane. The distance between the inner and outer membrane on each side was then calculated for points up to 500 nm from the midcell in both directions.

## Acknowledgments

We thank Martin Pilhofer for sharing the electron cryotomogram of an *E. coli* cell shown in Figure 4, Debnath Ghosal for helpful discussions, and Andrew Jewett for assisting with membrane-tracing software. This work was supported by the National Institutes of Health (grant R35 GM122588 to G.J.J.).

